# Hair follicle stem cell fate supports distinct clinical endotypes in *Hidradenitis Suppurativa*

**DOI:** 10.1101/2025.05.28.656362

**Authors:** Audrey Onfroy, Francette Jean-Louis, Philippe Le Corvoisier, Fanny Coulpier, Kevin Muret, Éric Bonnet, Raphaele Arrouasse, Caroline Boucle, Salwa Abid, Émilie Sbidian, Christina Bergqvist, Pierre Wolkenstein, Véronique Godot, Jean-François Deleuze, Yves Lévy, Piotr Topilko, Étienne Audureau, Sophie Hüe

## Abstract

Hidradenitis suppurativa (HS) is a severe skin disorder affecting 1% of the global population, with a complex and poorly understood pathogenesis involving aberrant keratinization and autoinflammation. It remains unclear whether autoinflammatory events precede or follow hyperkeratotic changes in hair follicle (HF) epithelia.

Using single-cell RNA sequencing, we characterized HF cell populations in HS patients and investigated their role in disease pathogenesis. We uncovered two distinct differentiation trajectories of HF stem cells (HF-SCs): one leading to interfollicular epidermis (IFE) basal cells enriched in inflammatory pathways, and another giving rise to outer root sheath (ORS) cells associated with keratinization. In HS lesions, both populations displayed altered inflammatory phenotypes and were closely linked to immune cell infiltration, pointing to a role in disease heterogeneity. By integrating clinical features with HF cell composition from 49 HS patients, we identified three major endotypes: (i) an inflammatory subtype, marked by T cell infiltration and an expansion of IFE basal cells; (ii) a keratinizing subtype, characterized by ORS enrichment and minimal inflammation; and a mixed subtype, exhibiting features of follicular remodeling, fistula formation, and variable immune involvement.

These findings provide novel insights into the epithelial-immune interactions that drive HS and support a stratified therapeutic approach tailored to the specific HF dysfunctions of each patient subgroup.

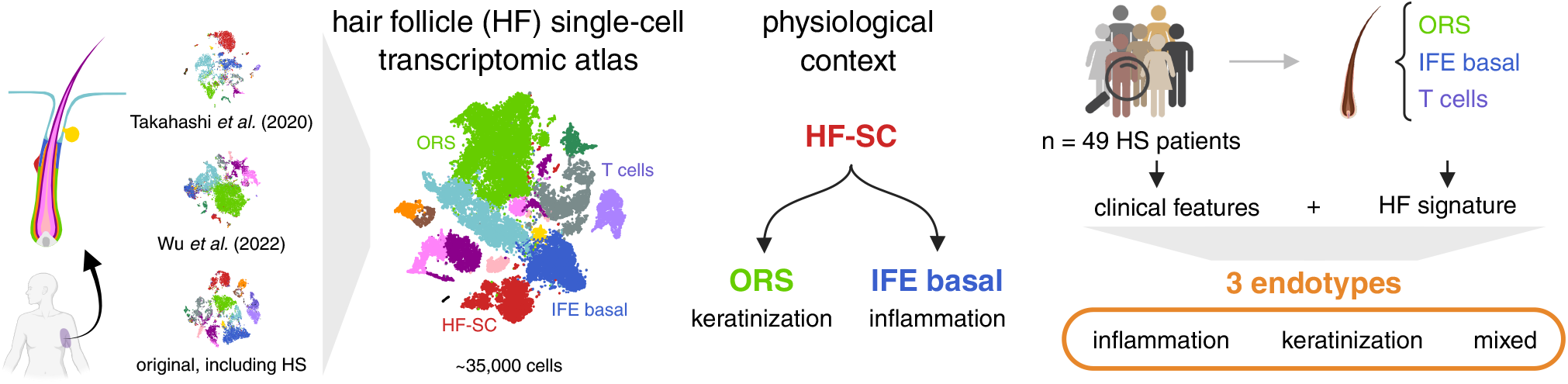

## Introduction

Hidradenitis suppurativa (HS) is a chronic, severe, inflammatory skin disorder with a prevalence of around 1% exerting a profound impact on patients’ quality of life (1). Characteristic lesions such as inflammatory nodules, abscesses, and sinus tracts develop in the axillae, inguinal, and gluteal areas, typically during or after puberty. HS is a complex disease with contributory of genetic, epigenetic hormonal, mechanical, microbial and lifestyle factors such as obesity and smoking. Current therapeutic options remain limited, and treatment outcomes are often unsatisfactory, largely due to an incomplete understanding of HS pathogenesis (2). In addition, pronounced disease heterogeneity has been increasingly recognized as a major factor influencing clinical response (3). The existence of distinct molecular and clinical endotypes may contribute to the failure of uniform therapeutic strategies, highlighting the urgent need for personalized approaches.

HS is recognized as an autoinflammatory keratinization disease because the pathogenic mechanisms of HS are intimately associated with aberrant keratinization and autoinflammation (4). In unaffected skin from predilection sites of patients with HS, the infundibular outer root sheath (ORS) shows hyperplasia and hyperkeratosis likely driven by an intrinsic dysregulation of the hair follicle stem cell (HF-SC) compartment. Notably, ORS cells exhibit replication stress, activating type I interferon responses (5). These epithelial alterations, in conjunction with external factors such as obesity and smoking, contribute to follicle occlusion and the accumulation of keratin and bacteria within the dilated hair follicle. The subsequent formation and rupture of cysts induce acute inflammation, characterized by an immune infiltrate of neutrophils, macrophages, dendritic cells, and T and B cells and increased expression of proinflammatory cytokines including IL1*β*, IL17, and TNF*α* (6). Treatment targeting inflammation, such as adalimumab, achieved clinical response in only 40% to 60% of patients, and newer biological therapies have not significantly exceeded this response rate (7). A major barrier to identifying novel targets and achieving successful clinical translation is the lack of understanding of the interplay between keratinization and inflammation.

In this study, we sought to better define the epithelial dynamics underpinning HS pathogenesis. Using single-cell RNA sequencing (scRNA-Seq), we characterized human hair follicle (HF) cell populations and uncovered two distinct differentiation trajectories of HF-SCs: one leading to interfollicular epidermal (IFE) basal cells enriched in inflammatory pathways, and the other to outer root sheath (ORS) cells, associated with keratinization. In HS patients, we observed that HF-SCs, IFE basal cells, and ORS cells exhibited a proinflammatory transcriptional profile and were closely associated with immune cell infiltration. Based on these observations, we hypothesized that the trajectory of HF-SC differentiation toward either lineage may influence disease heterogeneity. To investigate this, we analyzed HF cell composition in a cohort of 49 HS patients and identified three major endotypes: (i) an inflammatory endotype, marked by T cell infiltration and expansion of IFE basal cells; (ii) a keratinizing endotype, enriched in ORS with minimal inflammation; and (iii) a mixed endotype, featuring follicular remodeling, fistula formation, and variable levels of inflammation.

Our findings highlight the existence of distinct pathogenic mechanisms underlying HS-primarily inflammation, keratinization, or a combination of both-each rooted in specific HF-SC trajectories. This mechanistic stratification provides a new framework for understanding clinical heterogeneity in HS and opens avenues for personalized therapeutic strategies tailored to the dominant epithelial dysfunction within each endotype.

## Results

### Defining human hair follicle cell populations via integrated scRNA-Seq analysis

Since hair follicles play a central role in HS pathology, we decided to perform scRNA-Seq on HF cells. Skin samples were obtained from the perilesional lesions of five HS patients and compared to two healthy donors (HD) (see Table S1 for patients’ information). A scRNA-Seq analysis was conducted on HF cells, resulting in the acquisition of 12,111 cells that met the quality control criteria. The cells were annotated based on established HF markers and categorized into three main groups: immune cells, matrix cells and non-matrix cells (Figure 1A). The immune cells included *CD4*+ and *CD8*+ T cells, Langerhans cells, macrophages, and B cells.

**Fig. 1.**
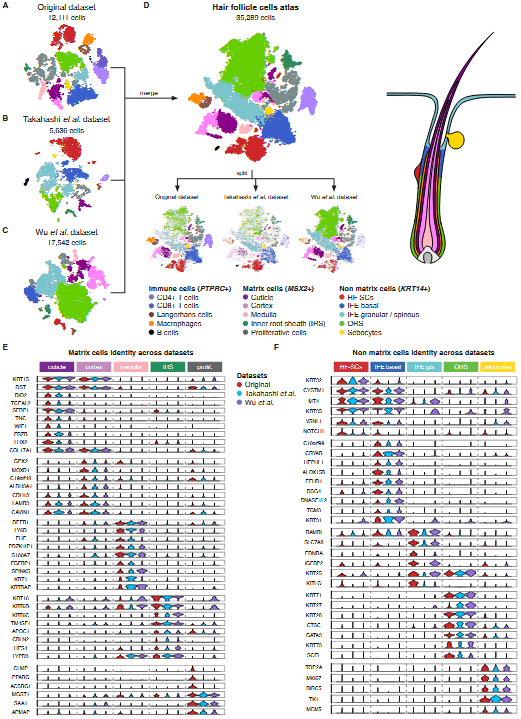
Transcriptomic atlas of hair follicle cells. **(A-C)** tSNE plots showing three individual datasets: our dataset (A), Wu *et al.* dataset (B) and Takahashi *et al.* dataset (C), colored according to cell type annotation. **(D)** After merging, the combined dataset contains 35,259 cells, colored according to cell type annotation. Split view shows the overlapping for all cell types across datasets. **(E-F)** Violin plots within matrix (E) or non-matrix (F) cells. Each panel is divided into five columns, assessing the identity of the cell type indicated on the top. Each column depicts three violin plots, associated with each dataset, showing the conserved expression of genes across dataset and laboratory. The legend key is shared between all panels.

To refine cell type annotation, we incorporated publicly available scRNA-Seq data from healthy individuals (8, 9) into a joint analysis with our dataset (Figure 1B-C). By merging these datasets, we constructed a comprehensive atlas of HF cells (Figure 1D). This integrative approach enabled the identification of shared gene signatures across datasets, thereby enhancing the resolution and categorization of HF cell populations. The matrix category was further divided into five populations based on their transcriptional profiles: i) cuticle (*KRT32, KRT35*), ii) cortex (*KRT31*), iii) medulla (*BAMBI*), iv) inner root sheath (IRS) (*KRT71, KRT28*) and v) proliferative cells, expressing high levels of *TOP2A* and *MKI67* genes (Figure 1E). The non-matrix category also regroups five populations: i) hair follicle stem cells (HF-SCs) expressing high levels of *KRT15, COL17A1, DIO2* and *TCEAL2*, ii)basal interfollicular epidermis (IFE) expressing *KRT15* and *COL17A1* but lacking *DIO2* and *TCEAL2* expression iii) granular and spinous IFE expressing *SPINK5* and *LY6D*, iv) outer root sheath cells (ORS) expressing cells expressing *KRT16, KRT6B* and *KRT6C*, and (v) sebocytes expressing *CLMP* and *PPARG* (Figure 1F).

This comprehensive molecular annotation revealed 15 transcriptionally distinct HF cell populations, consistently observed in both HS patients and healthy donors, irrespective of sex or sampling site (Figure 1). These findings provide a high-resolution view of HF cellular diversity.

### HF-SC fate mapping reveals two distinct lineages: ORS and IFE basal with unique molecular signatures

To investigate the differentiation dynamics of HF stem cells (HF-SCs), we constructed a diffusion map based on integrated scRNA-Seq data from both healthy donors and HS patients. This analysis revealed two conserved differentiation trajectories: one toward IFE basal and the other toward ORS (Figure 2A). These two distinct branches were consistently observed in both HS and healthy skin samples, indicating that this bifurcation represents a physiological feature of human HF-SC differentiation (Figure 2B). Trajectory inference using the Slingshot package confirmed the existence of these two lineages, which was further validated using the TInGa algorithm (Figure 2C). Along the IFE branch, cells continued to differentiate into granular and spinous layers of the epidermis.

**Fig. 2.**
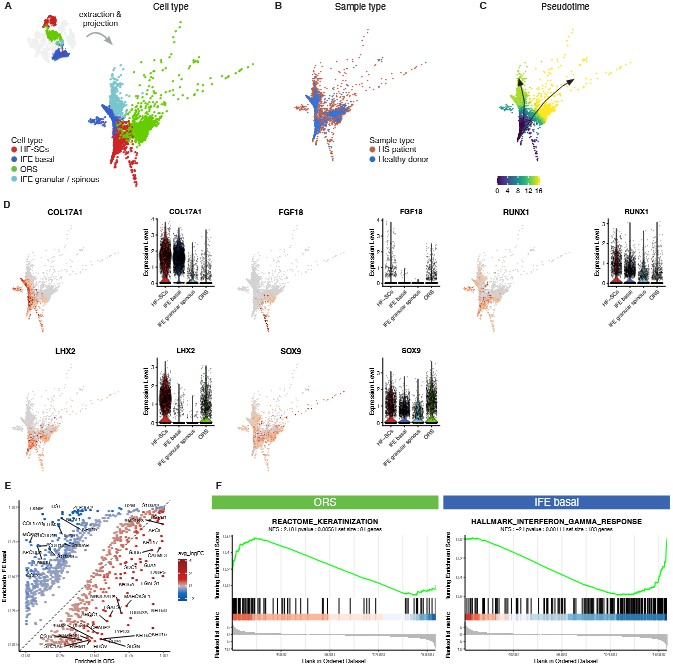
Non-matrix cells from HF-SCs follow two transcriptionally distinct lineages. **(A-C)** Diffusion map of merged HF-SCs, IFE basal, IFE granular/spinous and ORS from our dataset (Figure 1), illustrated in the top left corner. Cells are coloured based on cell type annotation (A), sample type (B) or pseudotime (C). **(D)** Feature plots and violin plots showing expression levels of selected genes. Violin plots are split by cell types. **(E)** Representation of differentially expression genes between ORS and IFE basal from all samples. X-and Y-axis respectively correspond to the proportion of cells within ORS and IFE basal cells with positive expression of the gene. Dashed line corresponds to genes expressed in same proportion of cells in ORS and IFE basal cells. One dot, corresponding to one gene, is coloured according to average log fold change (avg_logFC) between ORS and IFE basal cells. **(F)** GSEA plots made from average fold change of all genes, between ORS and IFE basal cells.

HF-SCs expressed a core set of stemness-associated genes, including key transcription factors such as SOX9, LHX2, and RUNX1, which help maintain stem cell quiescence and an undifferentiated state. Interestingly, their differentiated progeny retained subsets of these markers: IFE basal cells maintained *SOX9, RUNX1*, and *COL17A1*, while ORS cells retained *SOX9, LHX2*, and *FGF18* (Figure 2D).

Transcriptomic profiling revealed striking molecular differences between these two epithelial lineages (Figure 2E). Gene Set Enrichment Analysis (GSEA) showed enrichment of keratinization-related genes in the ORS branch, while IFE basal cells upregulated interferon response genes (Figure 2F). These lineage-specific gene expression patterns were reproducible across the two additional datasets (Figure S2). Al-together, our results suggest that HF-SCs differentiate into two functionally distinct lineages: ORS cells contribute to structural processes like keratinization, while IFE basal cells may play a more prominent role in mediating inflammatory responses.

### Inflammatory shift in hair follicle epithelial lineages in HS

Differential expression analysis of HF-SCs revealed fundamental transcriptomic changes between healthy donors and HS patients (Figure 3A). In HS patients, HF-SCs displayed an inflammatory phenotype, as evidenced by an enrichment in HLA class I molecule, *MIF* and *IFITM3* (Figure 3A). Furthermore, HF-SCs from HS patients were enriched for ribosomal genes suggesting a primed or activated transcriptional state. In contrast, HF-SCs from healthy donors showed higher expression of genes involved in DNA repair, such as *DDIT4*, and apoptosis pathway suggesting a more quiescent, homeostatic state (Figure 3A).

**Fig. 3.**
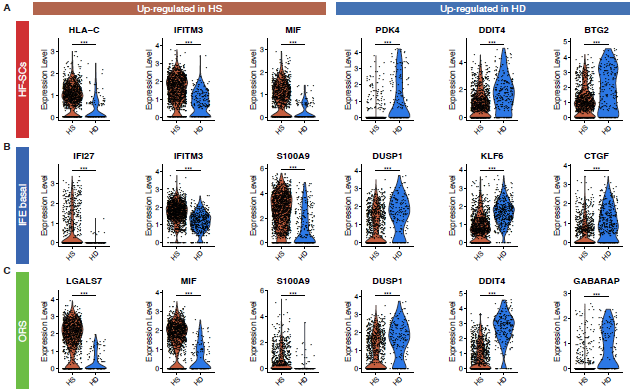
Non matrix cells show enrichment in inflammation markers in HS patients. Violin plots showing the expression level of differentially expressed genes between HS and HD, split by sample type, only within HF-SCs (A), IFE basal (B) or ORS (C). Statistical t-test p-value, ns: > 0.05, *: < 0.05 and > 0.01, **: < 0.01 and > 0.001 and ***: < 0.001.

A similar shift towards inflammation was observed in IFE basal cells from HS patients, which exhibited increased expression of inflammatory mediators such as *CCL2* and *S100A9*, along with a type I interferon signature (*IFI27, IFITM3*) (Figure 3B). Conversely, genes involved in cell cycle regulation (*DUSP1, KLF6*) and epithelial barrier integrity (*CTGF, CLDN1*) were downregulated in HS patients, potentially contributing to impaired epithelial homeostasis (Figure 3B).

In ORS cells, we likewise detected a shift toward inflammation and activation in HS patients. ORS cells overexpressed *MIF* and *S100A9*, consistent with immune activation (Figure 3C). Notably, we observed upregulation of *LGALS7* (galectin-7), a lectin predominantly expressed in stratified epithelia (10), and *ARF5*, a small GTPase involved in splicing, ribosome synthesis, and intracellular trafficking (11) (Figure 3C). These changes suggest an altered differentiation state of ORS cells in HS. By contrast, ORS cells from healthy donors expressed higher levels of *DUSP1* and *DDIT4*, genes crucial for cell cycle control and cellular stress response.

Together, these results indicate that HF-SCs, IFE basal, and ORS cells in HS patients adopt a transcriptional program characterized by inflammation, cellular activation, and impaired cell cycle and adhesion processes-highlighting broad epithelial dysregulation in HS skin.

### Cytotoxic and inflammatory immune cell signatures emerge around hair follicles in HS

We next investigated the immune cell populations present surrounding hair follicles. While healthy individuals exhibited minimal immune cell presence, HS patients displayed a marked infiltration of myeloid cells and/or T cells, accumulating around the hair follicles in the perilesional area (Figure 4A). To further investigate gene expression levels among immune cells, we generate a tSNE containing only immune cells from our dataset (Figure 4B).

**Fig. 4.**
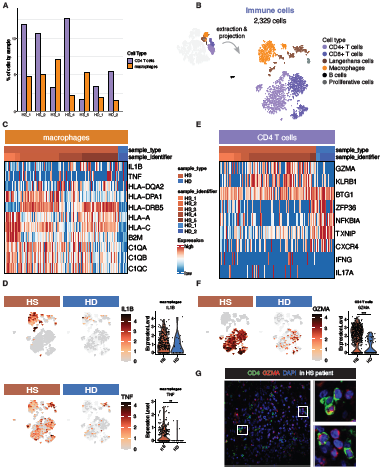
Immune cells in HS patients are activated compared to healthy donors. **(A)** Barplot showing the percentage of *CD4*+ T cells and macrophages in our dataset (Figure 1A), across sample identifiers. **(B)** tSNE plot showing 2,329 immune cells extracted from our dataset. Cells are colored based on cell type annotation. **(C)** Heatmap showing differentially expressed genes in macrophages, between HS and HD samples. **(D)** Feature plots and violin plots showing *IL1B* and *TNF* expression levels, split by sample type. On the tSNE, cells from complete immune cells dataset are shown as light background. The violin plots consider only macrophages. **(E)** Heatmap showing differentially expressed genes in *CD4*+ T cells, between HS and HD samples. Legend is shared with (C). **(F)** Feature plot and violin plot showing GZMA expression level, split by sample type. On the tSNE, cells from complete immune cells dataset are shown as light background. The violin plots consider only *CD4*+ T cells. **(G)** Section through hair follicles from a HS patient, immunolabeled from CD4, GZMA and DAPI. High magnification areas are denoted by square and represented on the right. Statistical t-test p-value, *: < 0.05, **: < 0.01 and ***: < 0.001. For additional features in immune cells, refer to Figure S3.

Differential gene expression analysis revealed upregulation of MHC class I and II genes, as well as components of the complement pathway, in macrophages from HS patients compared to healthy donors (Figure 4C). Notably, expression of *IL1B and* TNF levels was significantly elevated in HS macrophages (p = 0.058; p < 0.001, respectively) (Figure 4D), while *IL6* expression was undetectable (Figure S3A). These findings suggest the presence of a pro-inflammatory macrophage phenotype in close proximity to hair follicles within perilesional regions.

*CD4*+ T cells exhibited a cytotoxic phenotype, characterized by elevated expression of *GZMA* and *KLRB1*, which was confirmed by immunofluorescence (Figure 4E-G). While nearly all *CD4*+ T cells expressed *GZMA, IFNG* and *IL17* expression varied markedly between individuals (Figure 4E). TCR*α* analysis showed no evidence for invariant TCR usage, indicating that *CD4*+ T cell infiltration is polyclonal (Figure S3E).

Furthermore, CD8+ T cells in HS patients exhibited significantly higher expression of *IFNG, GZMB*, and *PRF1* compared to healthy donors, consistent with cytotoxic activity (Figure S3B-D). The generation of such cytotoxic T cells requires type I interferon and IL12 signaling (12). Therefore, the presence of these cells further reinforces the pivotal role of interferon in HS pathogenesis, as we have previously demonstrated (5).

### Self-organizing map analysis of 49 HS patients reveals distinct clinical patterns linked to HF cell composition and pathophysiological mechanisms

Given the potential contribution of HF-SC lineage divergence to HS heterogeneity, we sought to determine whether the IFE basal and ORS compartments are differentially enriched across patients. We hypothesized that this imbalance could reflect two distinct pathological mechanisms: one dominated by keratinization, linked to ORS enrichment, and the other driven by inflammation, associated with IFE basal expansion. To investigate this, we analyzed samples from 49 HS patients enrolled in the Fol-Hydra cohort (Table 1). RNA was extracted from five freshly isolated hair follicles per patient and subjected to qPCR analysis to quantify markers of ORS, IFE basal, and T cells, enabling us to infer the HF cellular composition in lesional skin. In parallel, and considering the central role of nicastrin (*NCSTN*) in maintaining skin homeostasis-as well as the fact that approximately 37% of known HS-associated variants map to this gene-we performed single nucleotide variant analysis of *NCSTN* in the same cohort. Notably, we identified two previously unreported variants, p.Leu30 and p.Gln83, both predicted to generate truncated proteins with potentially deleterious effects on function.

**Table 1.** Clinical characteristic of the 49 HS patients from the Fol-HYDRA cohort.

| Characteristic | N | Cluster A<br>N=8<sup>1</sup> | Cluster B<br>N=6<sup>1</sup> | Cluster C<br>N=9<sup>1</sup> | Cluster D<br>N=10<sup>1</sup> | Cluster E<br>N=16<sup>1</sup> | p-value<sup>2</sup> |
| --- | --- | --- | --- | --- | --- | --- | --- |
| **Clustering features** |  |  |  |  |  |  |  |
| Age | 49 | 23.5 [20.5;25.3] | 24.0 [20.8;25.8] | 28.0 [26.0;35.0] | 44.0 [39.0;51.0] | 31.0 [23.0;40.8] | <0.001 |
| Gender, females | 49 | 5 (62.5%) | 4 (66.7%) | 2 (22.2%) | 6 (60.0%) | 12 (75.0%) | 0.15 |
| Age at HS diagnosis | 48 | 18.0 [14.0;22.8] | 20.0 [19.0;21.0] | 20.0 [18.0;26.0] | 34.0 [28.0;41.0] | 27.5 [22.0;36.0] | <0.001 |
| Age at symptoms onset | 49 | 14.0 [14.0;15.5] | 14.5 [14.0;15.8] | 17.0 [16.0;18.0] | 26.5 [25.0;30.0] | 21.0 [17.0;23.8] | <0.001 |
| HS familial history | 38 | 3 (42.9%) | 4 (66.7%) | 0 (0.0%) | 1 (12.5%) | 5 (55.6%) | 0.021 |
| Smoking | 35 | 1 (25.0%) | 2 (50.0%) | 5 (71.4%) | 7 (77.8%) | 9 (81.8%) | 0.28 |
| Body mass index (BMI) | 49 | 25.0 [21.0;29.8] | 27.5 [25.3;32.0] | 25.0 [25.0;27.0] | 30.5 [26.8;33.0] | 23.5 [21.0;30.5] | 0.27 |
| LC1 Phenotype | 49 | 7 (87.5%) | 4 (66.7%) | 3 (33.3%) | 10 (100.0%) | 11 (68.8%) | 0.015 |
| LC2 Phenotype | 49 | 1 (12.5%) | 2 (33.3%) | 4 (44.4%) | 0 (0.0%) | 3 (18.8%) | 0.13 |
| LC3 Phenotype | 49 | 0 (0.0%) | 2 (33.3%) | 4 (44.4%) | 2 (20.0%) | 2 (12.5%) | 0.15 |
| Pilonidal sinus | 49 | 0 (0.0%) | 2 (33.3%) | 6 (66.7%) | 1 (10.0%) | 3 (18.8%) | 0.012 |
| Inflammatory nodule | 49 | 6 (75.0%) | 6 (100.0%) | 6 (66.7%) | 9 (90.0%) | 1 (6.3%) | <0.001 |
| Non-inflammatory nodule | 49 | 5 (62.5%) | 3 (50.0%) | 6 (66.7%) | 6 (60.0%) | 8 (50.0%) | 0.93 |
| Abscess | 49 | 0 (0.0%) | 1 (16.7%) | 4 (44.4%) | 1 (10.0%) | 1 (6.3%) | 0.076 |
| Open comedone | 49 | 2 (25.0%) | 6 (100.0%) | 5 (55.6%) | 8 (80.0%) | 4 (25.0%) | 0.002 |
| Bridge scar | 49 | 2 (25.0%) | 1 (16.7%) | 6 (66.7%) | 4 (40.0%) | 0 (0.0%) | 0.002 |
| Cookie cutter scar | 49 | 3 (37.5%) | 4 (66.7%) | 3 (33.3%) | 6 (60.0%) | 5 (31.3%) | 0.44 |
| Ice-pick scar | 49 | 2 (25.0%) | 4 (66.7%) | 6 (66.7%) | 4 (40.0%) | 3 (18.8%) | 0.084 |
| Draining fistula | 49 | 0 (0.0%) | 0 (0.0%) | 5 (55.6%) | 6 (60.0%) | 1 (6.3%) | <0.001 |
| Non-draining fistula | 49 | 2 (25.0%) | 0 (0.0%) | 5 (55.6%) | 7 (70.0%) | 1 (6.3%) | 0.001 |
| Papule | 49 | 1 (12.5%) | 1 (16.7%) | 4 (44.4%) | 2 (20.0%) | 5 (31.3%) | 0.63 |
| Folliculitis | 49 | 0 (0.0%) | 1 (16.7%) | 7 (77.8%) | 0 (0.0%) | 5 (31.3%) | <0.001 |
| Continuous damage | 49 | 3 (37.5%) | 5 (83.3%) | 9 (100.0%) | 6 (60.0%) | 4 (25.0%) | 0.001 |
| HS-PGA score | 49 | 2.00 [1.75;2.25] | 3.00 [2.25;3.00] | 3.00 [3.00;3.00] | 2.00 [2.00;3.00] | 1.00 [0.00;1.00] | <0.001 |
| Dissecting cellulitis | 49 | 0 (0.0%) | 0 (0.0%) | 1 (11.1%) | 0 (0.0%) | 0 (0.0%) | 0.47 |
| Folliculitis decalvans | 49 | 0 (0.0%) | 0 (0.0%) | 2 (22.2%) | 0 (0.0%) | 0 (0.0%) | 0.067 |
| Keloidal acne | 49 | 1 (12.5%) | 0 (0.0%) | 0 (0.0%) | 0 (0.0%) | 0 (0.0%) | 0.29 |
| **Illustrative features** |  |  |  |  |  |  |  |
| NCSTN variants | 49 | 0 (0.0%) | 2 (33.3%) | 0 (0.0%) | 0 (0.0%) | 0 (0.0%) | 0.033 |
| ORS-specific gene | 49 | 164 [69.0;260] | 387 [294;584] | 364 [331;667] | 407 [315;535] | 416 [317;464] | 0.023 |
| IFE basal-specific gene | 49 | 61.1 [44.8;64.5] | 57.8 [47.5;136] | 65.2 [36.5;74.2] | 67.5 [57.9;89.4] | 30.5 [23.8;43.8] | 0.033 |
| T cells-specific gene | 49 | 4.18 [2.24;6.5] | 7.5 [2.80;12.5] | 2.29 [1.56;5.1] | 2.57 [1.86;4.63] | 2.06 [1.04;2.50] | 0.068 |

<sup>1</sup>Median [25%;75%]; n (%)
<sup>2</sup>Kruskal-Wallis rank sum test; Fisher's exact test

To address clinical heterogeneity, we first performed an unsupervised classification of patients based solely on clinical parameters, independently of qPCR results or *NCSTN* genotyping. Using self-organizing maps (SOMs), we clustered patients according to their clinical features, revealing polarized distributions and five distinct clinical clusters (Figure 5A-B). Cluster boundaries are indicated by solid black lines, and a summary of clinical characteristics is provided in Table 1.

**Fig. 5.**
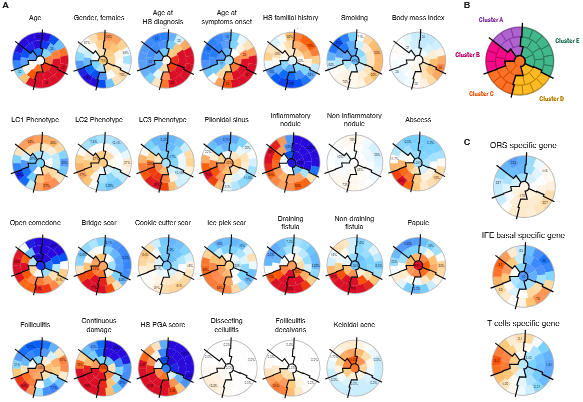
Clustering analysis of HS patients based on the Self-Organizing Maps (SOMs) methodology. **(A)** Unsupervised analysis by SOM placed all HS patients identified as globally similar for age, gender age at HS diagnosis, HS familial history and cigarette smoking within 1 of 40 small groupings (districts) throughout the maps. The more individuals resemble in terms of phenotyping, the closer they are placed on the map. Each individual map shows the mean values per district for each characteristic, blue colors indicating the lowest average values and red colors the highest, with detailed numbers written for a selection of representative districts in each SOM. **(B)** Close districts were combined to provide 5 suitable clusters of HS patients, labelled A to E. The clusters boundaries are delimited by solid black lines. **(C)** Representation of gene expression levels on the SOMs. These values were not taken into account while running the algorithm.

Patients in Cluster A (n = 8) and Cluster B (n = 6) comprised predominantly young, non-smoking women with early disease onset. The key difference between these clusters lies in their clinical presentation: in Cluster A, 7 out of 8 patients exhibited an LC1 phenotype, whereas all patients in Cluster B presented with open comedones (100%), ice-pick scars (66.6%) and continuous damage (83.3%). Additionally, 2 out of 6 patients in Cluster B had a pilonidal sinus, and notably, both patients with *NCSTN* variants belonged to this cluster. Strikingly, we observed a difference in HF composition, with an enrichment of IFE-associated gene expression in Cluster B. Both clusters A and B showed an infiltration of T cells (Figure 5A-B).

Cluster C (n = 9) included mostly male smokers, with early disease onset and no family history of HS. Only one patient exhibited an LC1 phenotype. Half of the patients presented with fistulas and continuous tissue damage, alongside the highest Hidradenitis Suppurativa Physician Global Assessment (HS-PGA) scores. This cluster showed co-enrichment of IFE basal and ORS gene expression but lacked T cell infiltration.

Cluster D (n = 10) consisted of the oldest patients, with the highest body mass index (BMI) and chronic disease features, including varied scar types and fistulas. Similar to Cluster C, they showed dual enrichment in IFE basal and ORS markers without T cell infiltration.

Cluster E (n = 16) comprised mostly women with an LC1 phenotype, but with distinct demographics: older age, history of smoking, normal BMI, and later disease onset. A significant proportion of patients had a family history of HS, and their nodules lacked inflammatory characteristics (Figure 5A-B). The HF cells in this cluster were enriched for ORS-associated gene expression, again without T cell infiltration (Table 1).

These results demonstrated a correlation between the sub-types identified by distinct clinical features and HF cells composition. In conclusion, these data suggest that the clinical features arise from three distinct pathophysiological processes: (i) a process in which inflammation is clearly predominant (Cluster A), (ii) a process in which keratinization occurs without inflammation (Cluster E), (iii) a process in which the two mechanisms are intertwined (Cluster B-C-D).

## Discussion

Deciphering the initial events that trigger follicular hyperplasia in HS remains particularly challenging due to the lack of suitable animal models. Unlike previous studies that have primarily focused on immune cell involvement, our research specifically investigates the epithelial compartments of the hair follicle, which we previously identified as key contributors to HS pathogenesis (5, 13). By integrating scRNA-Seq data from three independent datasets (8, 9), including our own, we generated a comprehensive atlas of HF cell populations. These populations were conserved across different anatomical sites and present in both healthy and inflamed skin, regardless of gender. However, discrepancies were noted in the classification of non-matrix cell populations-particularly between the dataset from Wu *et al.*, those from Takahashi *et al.* and our own. Notably, the population designated as inner bulge layer (IBL) by Wu and colleagues exhibited molecular features more consistent with ORS cells in our integrated analysis. These findings provide a high-resolution framework for understanding HF cellular diversity and dysfunction in HS.

Our study reveals that in HS patients, HF-SCs as well as basal IFE and ORS cells display an inflammatory phenotype that may act as an early trigger for immune cell recruitment. This leads to the infiltration of various immune populations within the HF niche. Among the five HS patients analyzed at single-cell resolution, only three exhibited significant T cell infiltration, highlighting the heterogeneity of immune responses and suggesting the existence of distinct inflammatory pathways. Once recruited, immune cells interact with HF-SCs, ORS, and IFE basal cells, further amplifying inflammation and disrupting tissue homeostasis. In some patients, cytotoxic T cells may contribute to epithelial damage through the secretion of GZMA, whereas in others, macrophage-derived TNF*α* and IL-1*β* may sustain chronic inflammation and impair HF-SC quiescence (14, 15), ultimately promoting aberrant tissue remodeling and follicular occlusion. The persistent activation of inflammatory pathways establishes a self-sustaining loop, exacerbating HS pathology and contributing to interpatient variability.

Given the central role of HF-SCs in epithelial homeostasis, we further explored their differentiation trajectories. Our analysis revealed that HF-SCs give rise to two distinct lineages: ORS cells and IFE basal cells, underscoring their dual contribution to epidermal renewal. Disruptions in HF-SC homeostasis may alter this differentiation balance, potentially skewing cell fate toward one or the other lineage. A bias toward IFE basal differentiation may contribute to epidermal hyperkeratinization and impaired barrier function, while a shift toward ORS differentiation may potentiate inflammatory signaling. This lineage-specific divergence may underlie aspects of HS clinical heterogeneity, including differential severity and treatment response.

Recognizing the heterogeneity within immune populations and HF-SC fate, we investigated whether these differences correlate with clinical heterogeneity. To address this, we conducted a comprehensive analysis of T cells, IFE basal cells, and ORS cells in the hair follicles of 49 HS patients. We identified five distinct HS endotypes based on clinical characteristics and demonstrated that the cellular composition of hair follicles varies accordingly. The inflammatory endotype (cluster A) was composed primarily of young, non-smoking women with high BMI and inflamed nodules. Their HF profiles were enriched in IFE basal cells and T cells, evoking a predominant inflammatory driver. This cluster aligns with the “regular phenotype” described by Van Der Zee (16), the LC1 pattern by Canoui-Poitrine (3), and Martorell’s inflammatory pattern (17). In contrast, the keratinizing endotype (Cluster E) was associated with normal BMI and smoking, with minimal inflammation. Hair follicles in this group were enriched in ORS cells, suggesting keratinization as the primary driver, similar to Martorell’s follicular subtype (17). The mixed cluster included Clusters B, C, and D, displaying substantial clinical heterogeneity. Clusters C and D shared features such as fistulae, continuous lesions, and enrichment in both IFE basal and ORS cells, but lacked T cell infiltration. Cluster C comprised mostly young, smoking men with early-onset disease, while Cluster D included older women with high BMI and later disease onset. Cluster B was distinct showing strong T cell infiltration, along with expanded IFE and ORS populations, and includes young women with early-onset disease but without fistulae or continuous involvement. It cannot be ruled out that Group D represents women from Group B at a more advanced stage of the disease. These findings indicate that follicular hyperplasia in HS may arise through distinct mechanisms-either inflammation, keratinization, or a combination of both. This highlights the necessity of implementing tailored therapeutic approaches to optimize patient outcomes.

While our study provides valuable insights, some questions remain unanswered. The factors determining whether a patient exhibits enrichment in ORS versus IFE basal cells require further investigation. Additionally, the substantial clinical heterogeneity observed among the clusters makes it unlikely that inflammation precedes keratinization, or vice versa. Longitudinal studies would be instrumental in determining whether patients transition between phenotypes over time. Moreover, analyzing treatment responses based on pathophysiological profiles could provide critical information for refining therapeutic strategies.

In conclusion, our findings suggest that distinct pathophysiological mechanisms underlie HS development. The cellular composition of lesional hair follicles provides a promising basis for stratified therapeutic approaches, allowing for interventions tailored to the inflammatory, keratinizing, or mixed nature of the disease. This mechanistic stratification may serve as a critical tool in guiding future clinical trials and improving precision medicine in HS.

## Materials and Methods

### Sex as a biological variable

Our study examined male and female patients, and sex-dimorphic effects are reported.

### Ethic of human samples

This study was conducted on a cohort comprising men and women with HS. HS patients were recruited from the Fol-HYDRA study and enrolled from the dermatology department of the Henri Mondor hospital between June 2022 and December 2023. The Clinical Investigation Center performed skin biopsies. This monocentric, prospective study was conducted in accordance with the declaration of Helsinki and approved by the appropriate ethics committee (CPP North-West IV: 2021-A02352-39, 13/01/2022). All patients gave written informed consent before study enrolment.

For scRNA-Seq analysis, hair follicle samples were from discarded plastic surgery specimens from two healthy individuals and five HS patients (Table S1). This study was conducted in accordance with the declaration of Helsinki and was approved by the French institutional committee (reference N° 20.12.11.69413). The main inclusion criteria were 1) history of HS according to the diagnostic provided by the experimented investigator and 2) aged 18 years or older.

### Human HF separation

Human samples were digested in 1 U/mL Dispase (Stemcell Technologies) overnight at +4°C. Individual hair follicles were pulled out using forceps. The isolated hair follicles were digested in TrypLE Express 1X (Gibco) for 30 minutes at room temperature, washed in DMEM + 10% FBS, and filtered through a 100 µm cell strainer.

### Single-cell RNA sequencing

HF cells were sorted (BD InfluxTM Cell Sorter) using a nuclear labeling DRAQ7TM Far-Red Fluorescent Live-Cell Impermeant DNA Dye (Abcam, ref: ab109202). Cells were collected in 0.04% BSA-RPMI solution.

For each individual, 25,000 cells were loaded into one channel of the Chromium system using the V3.1 single-cell reagent kit (10X Genomics). Following capture and lysis, cDNAs were synthesized, and then amplified by PCR for 12 cycles. The amplified cDNAs were used to generate Illumina sequencing libraries that were sequenced on one flow cell Nextseq500 Illumina.

### Analysis of scRNA-Seq data

Sequencing files were processed using 10x Genomics Cell Ranger 3.1.0. Reads were mapped on the GRCh37 (hg19) transcriptome. Raw count matrices were processed up to the figures presented here using RStudio, with R Markdown to generate fully traceable notebooks. To ensure version stability, a Singularity container containing R version 3.6.3 and all packages of interest was developed and used to compile notebooks. HTML reports, including codes and figures, are available in a Github repository as an intuitive website (see the Code and Data Availability section).

Individual datasets, associated with a single patient sample, were analyzed individually from the raw count matrix using Seurat V3 package (18). Cells having less than 500 genes were filtered out. Doublet cells were removed using scDblFinder tool (19) and scds in hybrid mode (20). Then, cells having more than 20% of UMI related to mitochondrial genes or more than 50% related to ribosomal genes were filtered out. The UMI count matrix for remaining cells was normalized using LogNormalize method implemented in *Seurat::NormalizeData* function. Single cells were annotated for cell type using a modified version of *Seurat::AddModuleScore* and cell type-specific marker gene sets in Table S2. According to previous publications associated with scRNA-Seq from skin (8, 9, 21), they encompass CD4+ T cells, CD8+ T cells, Langerhans cells, macrophages, B cells, cuticle, cortex, medulla, IRS, proliferative cells, ORS, IFE, HF-SC, sebocytes and melanocytes. To smoothen eventual mis-annotation, single cell level annotation was grouped by cluster. Clustering was generated as follows. First, 3000 highly variable features (HVFs) were identified using *Seurat::FindVariableFeatures*. Then, we generated a principal component analysis (PCA) with 100 components, from those 3000 HVFs, using *Seurat::RunPCA*. Distances between cells were computed using top 20 components of the PCA, with *Seurat::FindNeighbors*. Finally, a clustering was generated by *Seurat::FindClusters* with resolution 2. The cluster annotation was defined as the most represented cell type by cluster.

To build the main “atlas” dataset (Figure 1A), melanocytes clusters were removed in individual datasets prior to combination. Dataset combination was made using the *base::merge* function. For the populations-specific datasets associated with non-matrix cells, IFE basal and ORS, and immune cells, clusters of interest were selected, in the atlas dataset, based on their annotation. Genes expressed in less than 5 cells were removed. Merged count matrices were normalized using LogNormalize method and a 100 components PCA was made using the scaled expression of 2000 HVFs. Sample-specific effect was removed using *harmony::RunHarmony* on the PCA (22). The tSNE was built using *Seurat::RunTSNE* function.

For the dataset containing non-matrix cells, a diffusion map was used to generate a projection representative of transcriptomic changes between cells, using the implementation from destiny package (23). Trajectory inference was made using two distinct methods: slingshot (24) and TInGa (25), using cells in HF-SCs population as trajectory root, and the harmonized PCA as input. Slingshot output 7 lineages, either with dead end within HF-SC population, or toward IFE basal cells and ORS cells. TInGa can infer branching trajectory, thus was used for visualization purpose. Default parameters, except max_nodes set to 10, were used. Differential expression was conducted using *Seurat::FindMarkers*. Heatmaps were made using ComplexHeatmap package (26). Functional enrichment analyses were conducted using msigdbr package for the gene sets database, and clusterProfiler package for the analyses (27, 28). Gene sets score were computed using the *Seurat::AddModuleScore* function. When comparing the expression levels of scores between two populations, t-test implemented in the *stats::t.test* function was used.

To validate our findings, the same processing steps were applied to the dataset from Wu *et al.* and Takahashi *et al.*. For the former, input data were the FASTQ files, downloaded at project number OEP002321 on biosino.org portal. For the latter, the analysis was conducted from the count matrices downloaded at the accession number GSE129611 on Gene Set Omnibus portal. The detailed parameter setting are accessible in the compiled HTML files obtained from the R Markdown notebooks (see the Code and Data Availability section).

### RNA extraction and Real time Quantitative PCR

RNA extraction was performed according to the manufacturer’s protocol (RNeasy Micro Kit, QIAGEN Inc.). RNA was converted to cDNA with QuantiTect reverse transcription Kit (QIAGEN). Quantitative PCRs were performed using the Brilliant II SYBR GREEN QPCR Master Mix kit (Agilent Technologies) for the expression of three genes specific for T cells, IFE basal cells and ORS cells. Expression of the gene OAZ1 was used as a reference and the relative levels of each gene were calculated using the 2ΔΔCT method.

### Whole Genome Sequencing

Whole-genome sequencing was performed at the Centre National de Recherche en Génomique Humaine. After a complete quality control, genomic DNA (1 µg) was used to prepare a library for whole genome sequencing, using the Illumina TruSeq DNA PCR-Free Library Preparation Kit (Illumina Inc., CA, USA), according to the manufacturer’s instructions. After quality control and normalization, qualified libraries have been sequenced on a NovaSeqX+ platform from Illumina (Illumina Inc., CA, USA), as paired-end 150 bp reads. Samples were pooled on a NovaSeqX+ 25B flowcell in order to reach an average sequencing depth of 30X. Sequence quality parameters were assessed throughout the sequencing run and standard bioinformatics analysis of sequencing data was based on the Illumina pipeline to generate FASTQ files for each sample.

### Variant calling, annotation and analysis

The individual .g.vcf files were obtained through an in-house pipeline using the following tools: bwa-mem (v. 2.2.1), PicardTools (v. 2.26.9), Samtools (v. 1.16), Sambamba (v. 0.8.1), GATK (v. 4.2.3.0), and bedtools (v. 2.30.0). The alignment was performed on the reference genome, GRCh38. The .g.vcf files were aggregated and then transformed into .vcf files using the GATK CombineGVCFs and GenotypeGVCFs modules. The variants were filtered according to the commonly used criteria: (1) for SNVs: QD < 2, QUAL < 30, SOR > 3, FS >

60, MQ < 40, MQRankSum < −12.5, and ReadPosRankSum < −8; (2) for Indels: QD < 2, QUAL < 30, FS > 200, and ReadPosRankSum < −20. We annotated our variants with the SnpEff suite (v. 4.3.t) and the SnpEff variant database (v. 5.1). For this work, only the MANE NCSTN transcript (ENST00000294785) biallelic variants with a strong impact (the most deleterious) have been retained. Variant allele frequencies are provided by gnomAD (v. 4.1).

### Clustering analysis of patients

Clustering analyses consisted of using the self-organizing maps (SOMs) algorithm developed by Kohonen. In a nutshell, the SOMs algorithm assigns each individual to a specific area on the map based on their characteristics, placing similar individuals in proximity and distinct ones in remote locations, thus allowing to draw visual comparisons of unique or overlapping patient characteristics and disease subtypes. The SOMs were constructed by applying the approach developed within the Numero package framework (29) for the R statistical platform 1) building the SOMs with statistical verification of the robustness of the contrasts observed by permutation tests and 2) determining suitable groupings based on the direct visualization of data patterns and key characteristics of the dataset.

For illustrative purposes, Gabriel’s biplots were plotted to project the patients along the principal components axes from mixed principal component analysis according to their individual characteristics, colouring patients according to their final diagnosis or cluster and thus allowing for direct visual assessment of the discriminative ability of each subgrouping. For descriptive statistics, categorical variables are expressed as proportions (%), quantitative variables as means (± standard deviation [SD]) or medians (interquartile range [IQR]), as appropriate. Biological parameters were log-transformed due to the non-normality of their distribution. We compared groups from clustering by means of one-way ANOVA or Kruskal-Wallis tests for continuous variables and chi-square tests or Fisher’s exact tests for categorical variables. A p-value < 0.05 was considered significant. Descriptive statistics and between-clusters comparisons were realized using Stata software 15.1 (StataCorp, Tx, USA), and R 3.4.3 (R Foundation, Vienna, Austria; pca2d, Numero packages) for clustering analyses and visualizations.

## Study Approval

This monocentric, prospective study of the FolHYDRA cohort was conducted in accordance with the declaration of Helsinki and approved by the appropriate ethics committee (CPP North-West IV: 2021-A02352-39, 13/01/2022). All patients gave written informed consent before study enrollment. The scRNA-Seq study was conducted in accordance with the declaration of Helsinki and was approved by the French institutional committee (reference N° 20.12.11.69413).

## Code and Data Availability

The single-cell RNA Sequencing data underlying this article are available in Gene Expression Omnibus and can be accessed with the accession number GSE273109. Their computational processing is accessible at https://github.com/audrey-onfroy/Onfroy_et_al_scRNASeq_HS_2025. The Singularity container to render the notebooks is accessible on Zenodo, with record ID 15512051 (https://zenodo.org/records/15512051).

The record also contains a table summarizing the version of all the R packages involved in the study.

## Author Contributions

Conceptualization (AO, YL, PT, EA, SH); Data Acquisition (FJL, PLC, FC, RA, CBo, SA, ES, CBe,JFD); Analysis (AO, FJL, FC, EA,KM,EB); Interpretation (AO, FJL, EA, SH); Visualization (AO, EA, SH); Writing original draft (AO, SH); Writing and review (AO, FJL, PLC, ES, PW, VG, PT, EA, KM, JFD, EB, SH); Project Administration (PW, VG, YL, PT, EA, SH); Funding Acquisition (SH).

All authors have declared that no conflict of interest exists.

## Acknowledgments

We are grateful to the IMRB genomic facility. We thank the staff from FACS for technical assistance.

## Supplementary Tables

**Supplementary Table 1.** Clinical characteristics of patients (n=7) for scRNA-Seq analysis. Main features of the 7 patients associated with the scRNA-Seq dataset. NA: not available

| Patient | Sex | Age (years) | Phenotype | Hurley Score | Sampling site |
| --- | --- | --- | --- | --- | --- |
| HS_1 | M | 44 | LC2/3 | III | axillary |
| HS_2 | M | 26 | LC1/3 | III | axillary |
| HS_3 | M | 28 | LC2 | II | pubis |
| HS_4 | F | 20 | LC1 | III | axillary |
| HS_5 | F | 21 | LC1 | III | axillary |
| HD_1 | M | 31 | NA | NA | pubis |
| HD_2 | F | NA | NA | NA | pubis |

**Supplementary Table 2.**
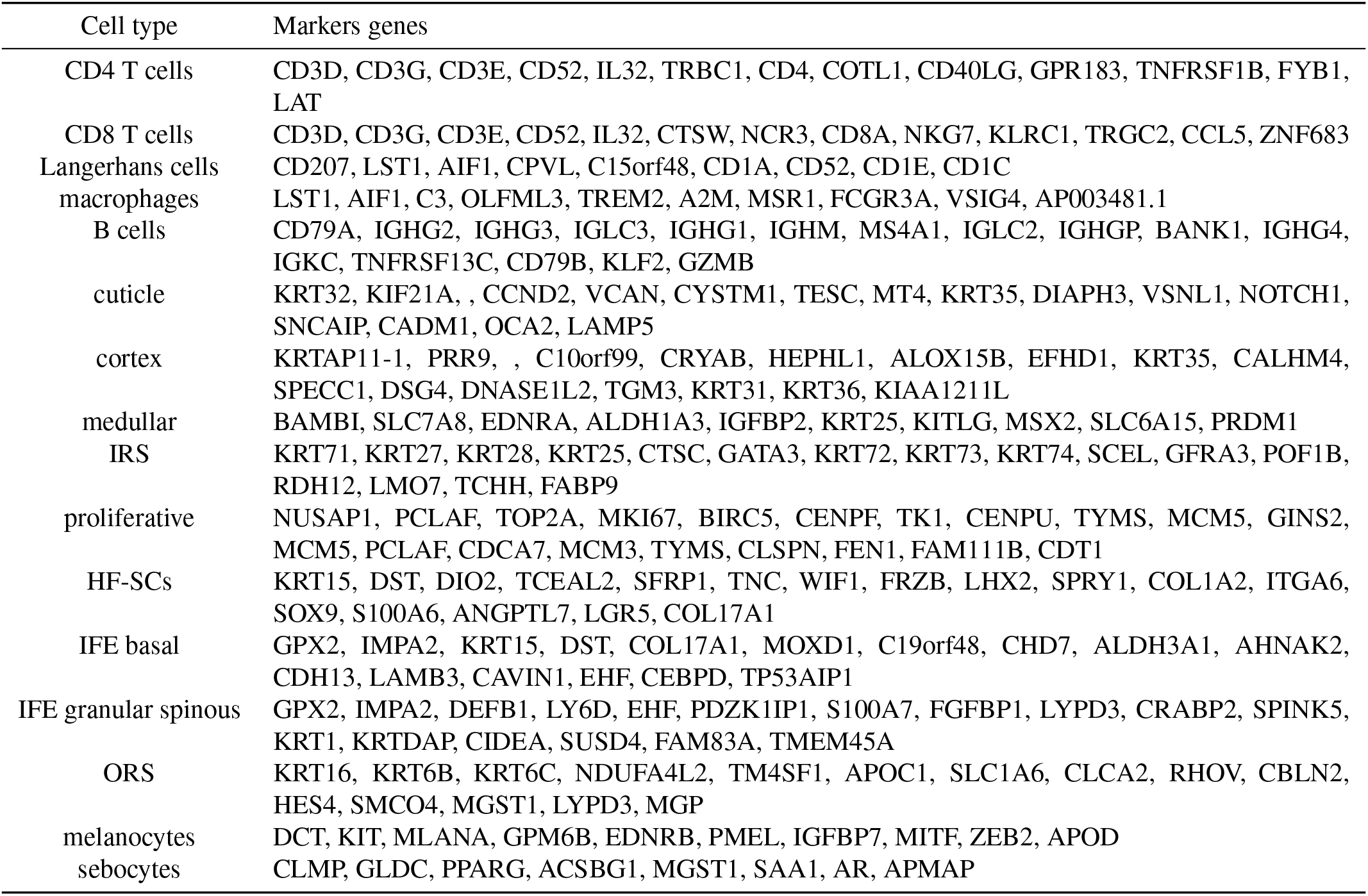
Gene markers for single-cell level cell type annotation of scRNA-Seq data. Each tab is named by a cell type. The tab content corresponds to the genes that have been used to annotated cell types in the scRNA-Seq data. This table is also provided, as R code, in the corresponding notebook (1_build_metadata.Rmd) on the Github repository.

## Supplementary Figures

**Supplementary Figure 1.**
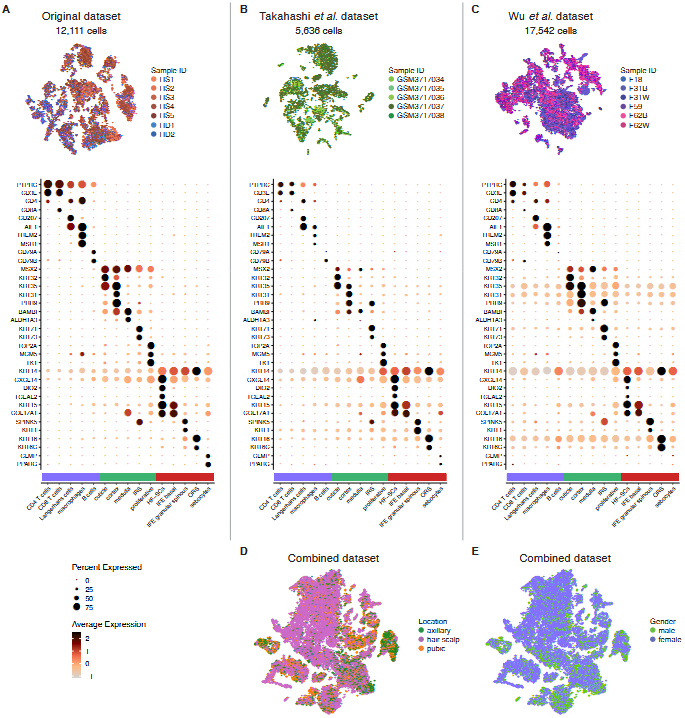
Additional features regarding scRNA-Seq datasets. **(A-C)** tSNE plot and dotplot made from our dataset A), Takahashi *et al.* dataset **(B)** and Wu *et al.* dataset **(C)**, colored according to sample identifier. The dot plot assesses cell type annotation in each dataset. Dot size represents the percentage of cells with positive expression for each marker. Dot color represents the scaled expression level of markers across all cells within each cell type. **(D-E)** tSNE plot of cells from the combined dataset, colored according to tissue of origin (D) or patient gender (E).

**Supplementary Figure 2.**
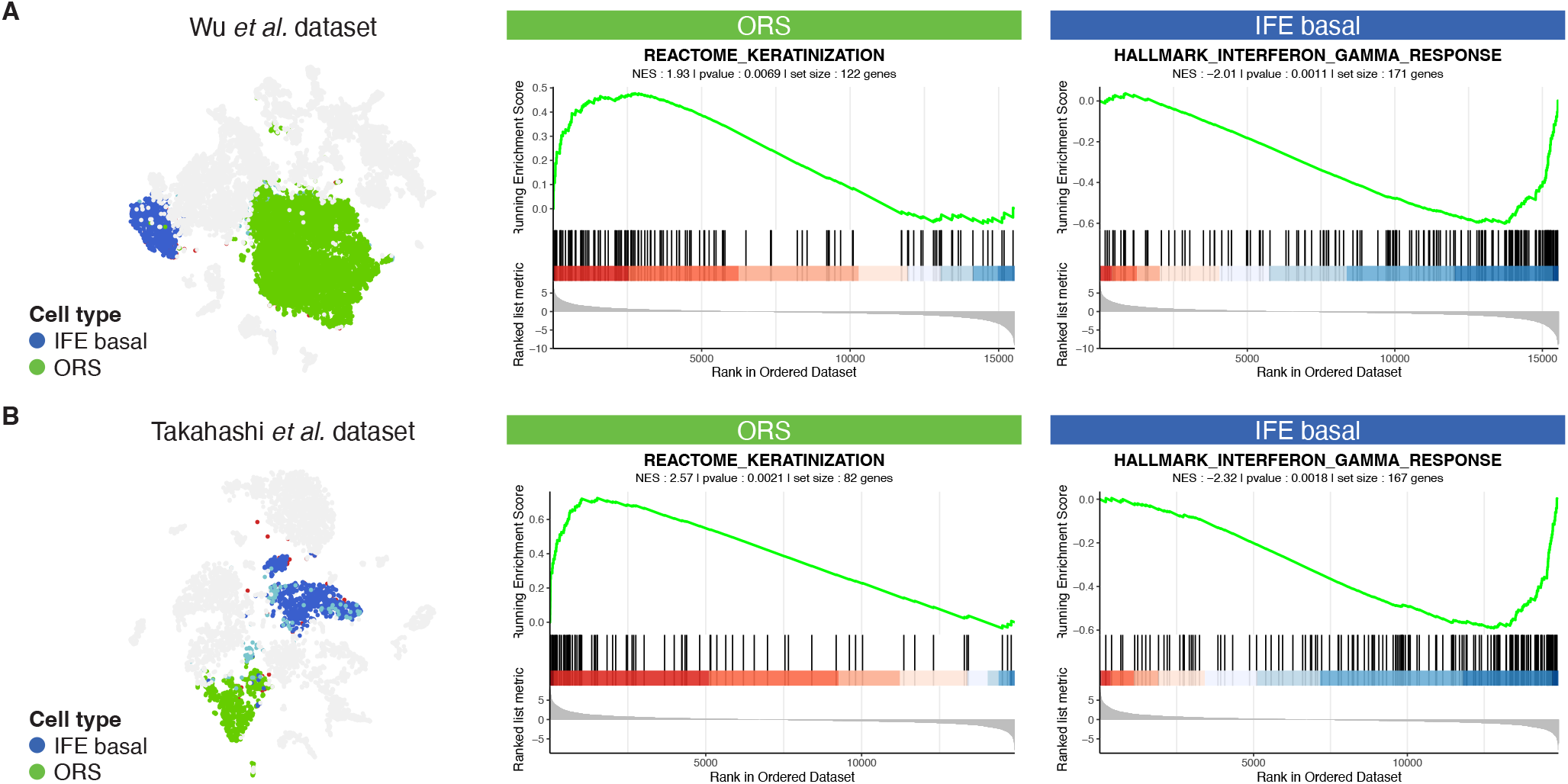
The activity of IFE basal and ORS is conserved across datasets. **(A-B)** tSNE highlighting IFE basal and ORS from the Wu *et al.* dataset (A) or the Takahashi *et al.* dataset (B). The associated GSEA plots are made from average fold change of all genes, between ORS and IFE basal cells, in the corresponding dataset.

**Supplementary Figure 3.**
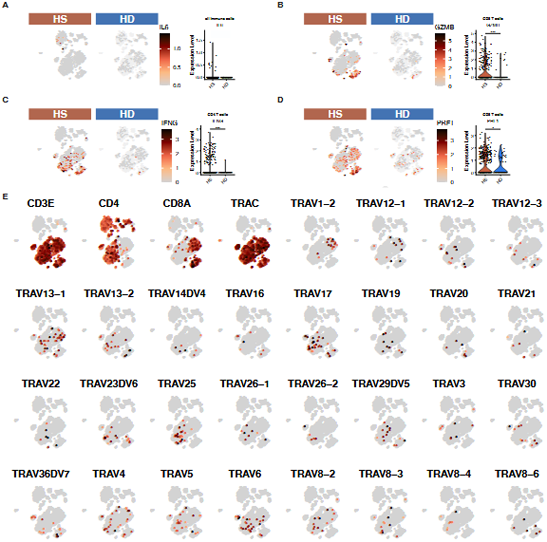
T cells features across sample types. **(A-D)** Feature plots and violin plots showing gene expression levels, split by sample type. On the tSNE, cells from complete immune cells dataset (Figure 4B) are shown as light background. The violin plots consider the cells population indicated in the title. Statistical t-test p-value, *: < 0.05, **: < 0.01 and ***: < 0.001. (E) Feature plots showing expression levels of genes related to T cells receptor alpha chain.

